# Na_v_1.5 expressed in TrkB^+^ sensory neurons mediates paclitaxel-induced mechanical pain hypersensitivity

**DOI:** 10.64898/2026.03.19.713015

**Authors:** Francisco Isaac F. Gomes-Aragão, Beatriz L. Adjafre, Sang Hoon Lee, Arthur Silveira Prudente, Gabriel V. Lucena-Silva, Conceição Elidianne Anibal Silva, Luiz Gustavo Kanada, Danilo Roman-Campos, José C. Alves-Filho, Fernando Q. Cunha, Stephen G. Waxman, Sulayman Dib-Hajj, Temugin Berta, Thiago M. Cunha

## Abstract

Chemotherapy-induced neuropathic pain (CINP) is a frequent and debilitating adverse effect of anti-tumor therapies, for which current treatments are largely non-specific and offer limited efficacy. Identifying molecular mechanisms that drive CINP may enable the development of targeted therapeutic strategies. Here, we demonstrate that paclitaxel-induced mechanical pain hypersensitivity in mice occurs independently of classical Na_v_1.8^+^ nociceptors but critically depends on TrkB^+^ sensory neurons. Transcriptomic analysis of TrkB^+^ sensory neurons revealed selective expression of *Scn5a*, which encodes the voltage-gated sodium channel Na_v_1.5, a channel classically associated with cardiac excitability. Importantly, *SCN5A* expression was also detected in human primary sensory neurons, indicating potential translational relevance. Functional studies further showed that *Scn5a* knockdown, using small interfering RNA, significantly attenuates paclitaxel-induced mechanical pain hypersensitivity. Together, these findings identify TrkB^+^ sensory neurons as key drivers of CINP and reveal Na_v_1.5 as a previously unrecognized contributor to chemotherapy-induced neuropathic pain. Targeting Na_v_1.5 in TrkB^+^ sensory neurons may therefore represent a novel therapeutic strategy for the treatment of CINP.

## Introduction

Neuropathic pain, including chemotherapy-induced peripheral neuropathy (CINP) caused by paclitaxel, is a common and debilitating condition that can limit cancer treatment efficacy, yet its underlying molecular mechanisms remain poorly understood (Seretny et al., 2014; Boyette-Davis et al., 2018; Fumagalli et al., 2021; Finnerup et al., 2021). Mechanical allodynia, a hallmark of neuropathic pain, arises from altered processing in mechanosensory pathways involving distinct dorsal root ganglion neurons (DRG) subtypes. Whereas Nav1.8+ primary sensory neurons seem to be dispensable for mechanical pain hypersensitivity triggered by peripheral nerve injury (Abrahamsen et al., 2008), low-threshold mechanoreceptors, particularly TrkB+ neurons, have been identified with a critical role, as they are both necessary and sufficient for the development of touch-evoked pain under neuropathic conditions (Dhandapani et al., 2018; Santana-Varela et al., 2021; Handler & Ginty, 2021). Apart from that, the relative contribution of Nav1.8+ nociceptors *versus* TrkB+ sensory neurons for paclitaxel-induced mechanical pain hypersensitivity has not been demonstrated. Herein, we report that Nav1.8⁺ sensory neurons were not involved in paclitaxel-induced mechanical hypersensitivity. On the other hand, chemogenetic silencing of TrkB⁺ sensory neurons reversed this process. Reanalysis of single-cell RNA-sequencing datasets from both mouse and human identified a selective expression of *Scn5a* (Nav1.5), a non-classical nociceptive voltage-gated sodium channel, in TrkB⁺ neurons, which was further validated experimentally and found to be upregulated after paclitaxel treatment. Finally, knockdown of Nav1.5 in primary sensory neurons attenuated paclitaxel-induced mechanical pain hypersensitivity, highlighting it as a key mediator of chemotherapy-induced neuropathic pain.

## Materials and methods

### Animals

All procedures were approved by the Ethics Committee for Animal Research of the University of São Paulo (1412/2025), the Institutional Animal Care and Use Committee (25-01-27-01) at the University of Cincinnati, following IASP and NIH guidelines, and reported according to ARRIVE (Percie Du Sert et al., 2020). C57BL/6J (Cat# 000664), Nav1.8^Cre^ (Cat# 036564), TrkB^CreER^ (Cat# 027214), ROSA-DTA^flox/flox^ (Cat# 009669), hM4Di^flox/flox^ (Cat# 026219), and Ai14^flox/flox^ (Cat# 007914) mice were obtained from Jackson Laboratory. Nav1.8^Cre/0^ mice were crossed with ROSA-DTA^flox/flox^ to deplete nociceptive neurons (Abrahamsen et al., 2008), while TrkB^CreER^ mice were crossed with Ai14^flox/flox^ and hM4Di^flox/flox^ to enable labelling and silencing of TrkB⁺ neurons, respectively. TrkB^CreER^ mouse lines received tamoxifen (200 µL, 1%, s.c.) for five consecutive days at 6–8 weeks of age (Santana-Varela et al., 2021).

### Paclitaxel-induced mechanical pain hypersensitivity

Mice received vehicle or paclitaxel (8 mg/kg, i.p.; ONTAX®, Libbs) on days 0, 2, 4, and 6 (Silva et al., 2022). Behavioral assessments included dynamic mechanical allodynia, evaluated by brush-evoked responses scored from 0 to 3, with median scores calculated (Cheng et al., 2017), as well as von Frey testing to determine mechanical thresholds (0.008–0.6 g) and paw withdrawal frequency to repeated 0.02 g stimulation (Decosterd and Woolf, 2000; Fonseca et al., 2020).

### siRNA Scn5a knockdown and behavioural testing

Intrathecal injections were performed by spinal puncture using a 30-gauge needle between the L5 and L6 levels, as described previously (PMID: 38888973). Mice received intrathecal injections of siRNAs targeting *Scn5a* (#s73373) at 2 µg per delivery in the transfection agent polyethylenimine. A non-targeting control siRNA (#4390846) was used as a negative control (all siRNA oligonucleotides were from Thermo Fisher Scientific).

### Chemogenetic inactivation of TrkB-positive sensory neurons

TrkB-^CreER^; hM4Di^flox/0^ mice and controls received paclitaxel and, on day 8, a single paw injection of clozapine N-oxide (20 nmol, 10 µL; Tocris, Cat# 6329) or vehicle, followed by assessment of mechanical pain threshold and withdrawal frequency as previously described (adapted from Saika et al., 2020).

### Analysis of publicly available scRNA-seq datasets

The pre-processed scRNA-seq datasets GSE139088 (Sharma et al., 2020) and GSE154659 (Renthal et al., 2020) were obtained from GEO, and the snRNA-seq dataset phs001158.v2.p1 (Tavares-Ferreira et al., 2022) from dbGaP. Analyses were performed in RStudio using Seurat (Hao et al., 2023).

Quality control was applied per dataset: Sharma et al.—cells with >5% mitochondrial RNA or <1,000 genes/counts were removed; Renthal et al.—cells with >10% mitochondrial RNA, >15,000 counts, or >5,000 genes were excluded; Tavares-Ferreira et al.—cells with >10% mitochondrial RNA, <6,000 genes, or <30,000 counts were excluded. Data were then library-size normalized, log1p-transformed, and scaled.

Highly variable genes were identified using the VST method, selecting the top 3,000 HVGs. PCA was performed using 45 components (Sharma) and 30 components (Renthal and Tavares-Ferreira). A shared nearest neighbour was constructed, followed by Louvain clustering (resolution: 0.8 for Sharma; 1.0 for Renthal and Tavares-Ferreira). Clusters were visualized using t-SNE or UMAP and explored with violin plots, bubble plots, and heatmaps.

### RNA scope in situ hybridization

DRGs (L3–L5) were collected, fixed in 4% PFA, cryoprotected in sucrose, embedded in OCT (Tissue-Tek), and sectioned (14–20 µm). RNA scope® was performed using the Multiplex Fluorescent Reagent kit v2 (Cat# 323110) with probes for *Scn5a* (Cat# 429881) and *Ntrk2* (Cat# 423611-C2). Sections were immunoassayed with anti-Neurofilament 200 (Sigma, Cat# N4142) and Alexa Fluor 594 secondary antibody (Invitrogen), counterstained with Nissl (Cat# N21480), and mounted with ProLong™ Gold (Cat# P36934). Neurons with >5 puncta were considered positive, and images were acquired (Keyence BZ-X810) and analysed using ImageJ (Schindelin et al., 2012).

### Quantitative Reverse-Transcription PCR (qRT-PCR)

Samples were stored in Trizol (Sigma) at −80 °C, and total RNA was extracted, reverse transcribed using the High-Capacity kit (MultiScribe®, Life-Invitrogen), and analysed by qRT-PCR (10 µL; 95 °C/40 cycles) using Gapdh, Scn5a, Scn10a, Th, Trpv1, Nefh, and Ntrk2 primers (Table 1). Reactions were run on QuantStudio™ 5 Real-Time PCR System, 96-well, 0.1 mL (ThermoFisher Scientific), with Gapdh as control, and data were analysed by the comparative 2^ΔΔCT method (Schmittgen & Livak, 2008).

### Human samples and Reverse-Transcription PCR (RT-PCR)

Lumbar DRG tissues were collected from a deidentified and non-diseased human donor. cDNA was synthesized from these tissues as described above, and RT-PCR analysis was performed with primers in Table S1.

### Statistical Analyses

Data were expressed as mean ± s.e.m. and analysed using two-way repeated measures ANOVA with Bonferroni’s post hoc test, or Student’s t-test and Friedman’s test when appropriate, with significance set at *p* < 0.05, using IBM SPSS Statistics for Windows 27.0.1.0 or GraphPad Prism version 9.0.2.161 for Windows.

## Results and Discussion

### 1. Paclitaxel-induced mechanical pain hypersensitivity does not require Nav1.8+ nociceptors

The role of Nav1.8⁺ sensory neurons in mechanical pain hypersensitivity is well established in inflammatory conditions but remains controversial in neuropathic pain (Daou et al., 2016; Abrahamsen et al., 2008). To test whether Nav1.8-lineage sensory neurons are required for paclitaxel-induced mechanical hypersensitivity, we used Nav1.8-Cre/DTA mice (Yang et a., 2023), in which Cre-dependent expression of diphtheria toxin fragment A (DTA) selectively ablates Nav1.8/*Scn10a*-expressing sensory neurons (that is, putatively most nociceptors, Fig. 1A). Notably, paclitaxel induced similar mechanical pain hypersensitivity in in Nav1.8-Cre/DTA mice and control mice (Fig. 1 B–D), arguing against an obligate role for Nav1.8⁺ nociceptors in this CINP phenotype. Consistent with effective nociceptor depletion, gene expression analysis showed reduced markers associated with nociceptor populations, including *Scn10a* (Nav1.8), *Trpv1*, and *Th* (Fig. 1E). In contrast, transcripts enriched in large-diameter, mechanoreceptor neuronal populations were preserved or enriched, including *Ntrk2* (TrkB) and *Nefh* (Fig. 1E). Together, these data suggest that paclitaxel-evoked mechanical allodynia can develop independently of Nav1.8⁺ nociceptors and instead may be driven by Nav1.8-negative sensory populations that remain intact in this model, including TrkB-positive mechanosensory neurons.

**Figure 1.**
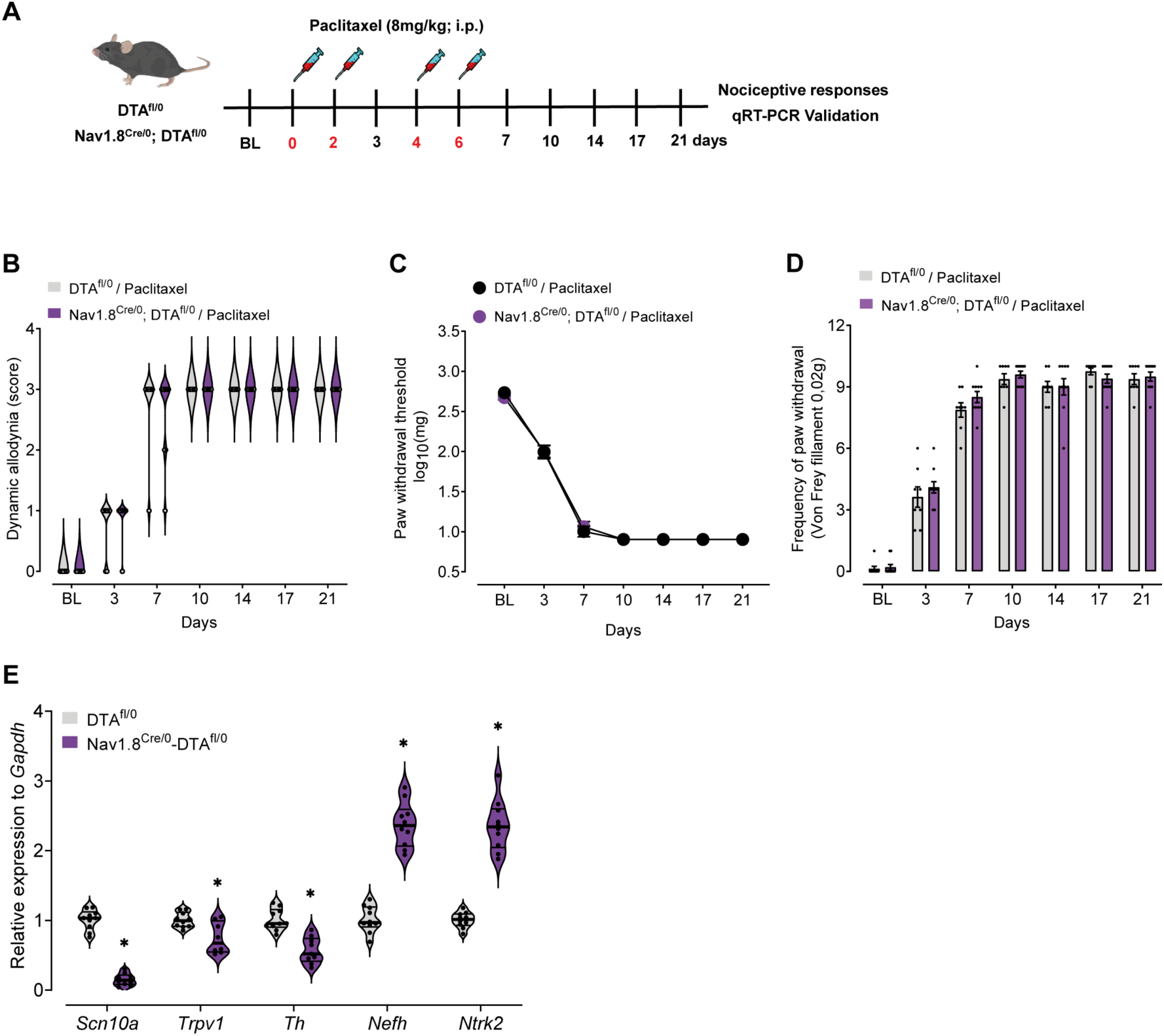
Role of Nav1.8+ sensory neurons in paclitaxel-induced mechanical pain hypersensitivity. **(A)** Schematic diagram of the time course of behavioral responses of mice devoid of nociceptors and controls treated with paclitaxel (8mg/kg; i.p). **(B)** Score of dynamic allodynia, Friedman Test (n= 8-10), **(C – D)** Mechanical paw withdrawal threshold and number of paw withdrawals to a 0.02 g von Frey filament, Two-way ANOVA, Bonferroni post-test (n= 8-10). **(E)** Molecular markers of cell populations in the DRG from control and mutant mice, p*<0.05 versus DTA^fl/0^, Student’s t Test, n = 10. *Scn10a*: Sodium Voltage-Gated Channel Alpha Subunit 10; *Trpv1*: Transient Receptor Potential Cation Channel Subfamily V Member 1; *Th*: Tyrosine Hydroxylase; *Nefh*: Neurofilament Heavy Chain; *Ntrk2*: Neurotrophic Receptor Tyrosine Kinase 2.

### 2. Silencing TrkB-positive sensory neurons attenuates paclitaxel-induced mechanical pain hypersensitivity

TrkB-positive sensory neurons have been implicated in mechanical pain hypersensitivity after peripheral nerve injury (Dhandapani et al., 2018; Santana-Varela et al., 2021). Since they are Na_v_1.8-negative fibers and their role in CINP has not been established, we investigated whether TrkB⁺ sensory neurons are functionally required for paclitaxel-induced mechanical pain hypersensitivity. To suppress TrkB⁺ neuron activity, we used an inhibitory DREADD (Designer Receptors Exclusively Activated by Designer Drugs) strategy (Zhu et al., 2016) in which TrkB⁺ neurons expressed the Gi-coupled hM4Di receptor, and clozapine-N-oxide (CNO) administration reversibly reduced excitability and neurotransmitter release. Mice expressing inhibitory DREADD receptors in TrkB-positive sensory neurons were subjected to the paclitaxel treatment, and mechanical sensitivity was assessed over time (Fig. 2A). CNO-mediated chemogenetic silencing of TrkB⁺ neurons produced a significant reduction in paclitaxel-induced mechanical pain hypersensitivity, while baseline responses remained unchanged (Fig. 2B-C). These findings indicate that TrkB⁺ sensory neurons play a critical and selective role in driving chemotherapy-induced mechanical pain hypersensitivity.

**Figure 2.**
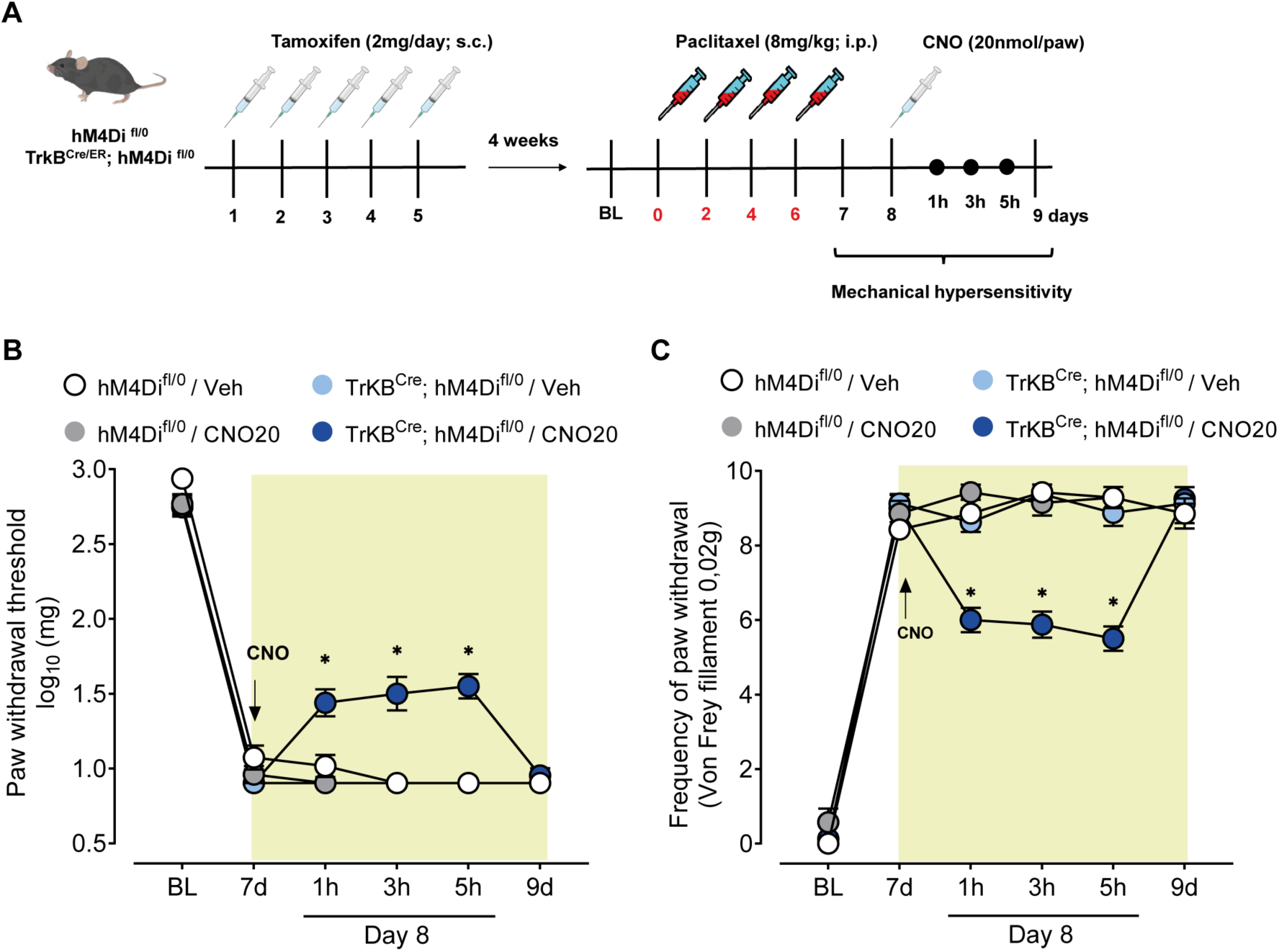
TrkB+ sensory neurons are involved in paclitaxel-induced mechanical pain hypersensitivity. **(A)** Schematic diagram of the time course of behavioral responses after chemogenetic silencing of TrkB+ sensory neurons after paclitaxel chemotherapy treatment (8mg/kg; i.p). (**B-C**) Mechanical pain hypersensitivity was evaluated in paclitaxel-treated mice before and after vehicle or clozapine N-oxide (CNO, 20 nmol/paw) to locally silence TrkB+ sensory neurons, *p<0.05 versus TrkB^CreER^ hM4Di^fl/0^ + Vehicle, 2-Way ANOVA, Bonferroni post-test, n =7-8.

### 3. Mice and human molecular profile of *TrkB+* sensory neurons revealed the expression of Nav1.5

Given the functional role of TrkB⁺ neurons in paclitaxel-induced mechanical pain hypersensitivity, we next sought to identify molecular effectors selectively expressed by this population that could be therapeutically tractable. Thus, we re-analyzed published mouse DRG single-cell RNA-seq data (GSE139088) and focused on TrkB⁺ sensory neurons subpopulation (Adelta LTMR fibers, Sharma et al., 2020). Strikingly, among the most enriched genes (top 25) in TrkB⁺ neurons, *Scn5a* (encoding Nav1.5), an non-classical nociceptive voltage gated sodium channels (Fig. 3A), emerged as one of the most selectively enriched transcripts within this cluster (Fig. 3A) and showed comparatively restricted expression across DRG neuronal subtypes (Fig. 3B). This observation is consistent with earlier reports that Nav1.5, although classically studied in the heart, is also present in a discrete subset of primary sensory neurons (Renganathan et al., 2002).

**Figure 3.**
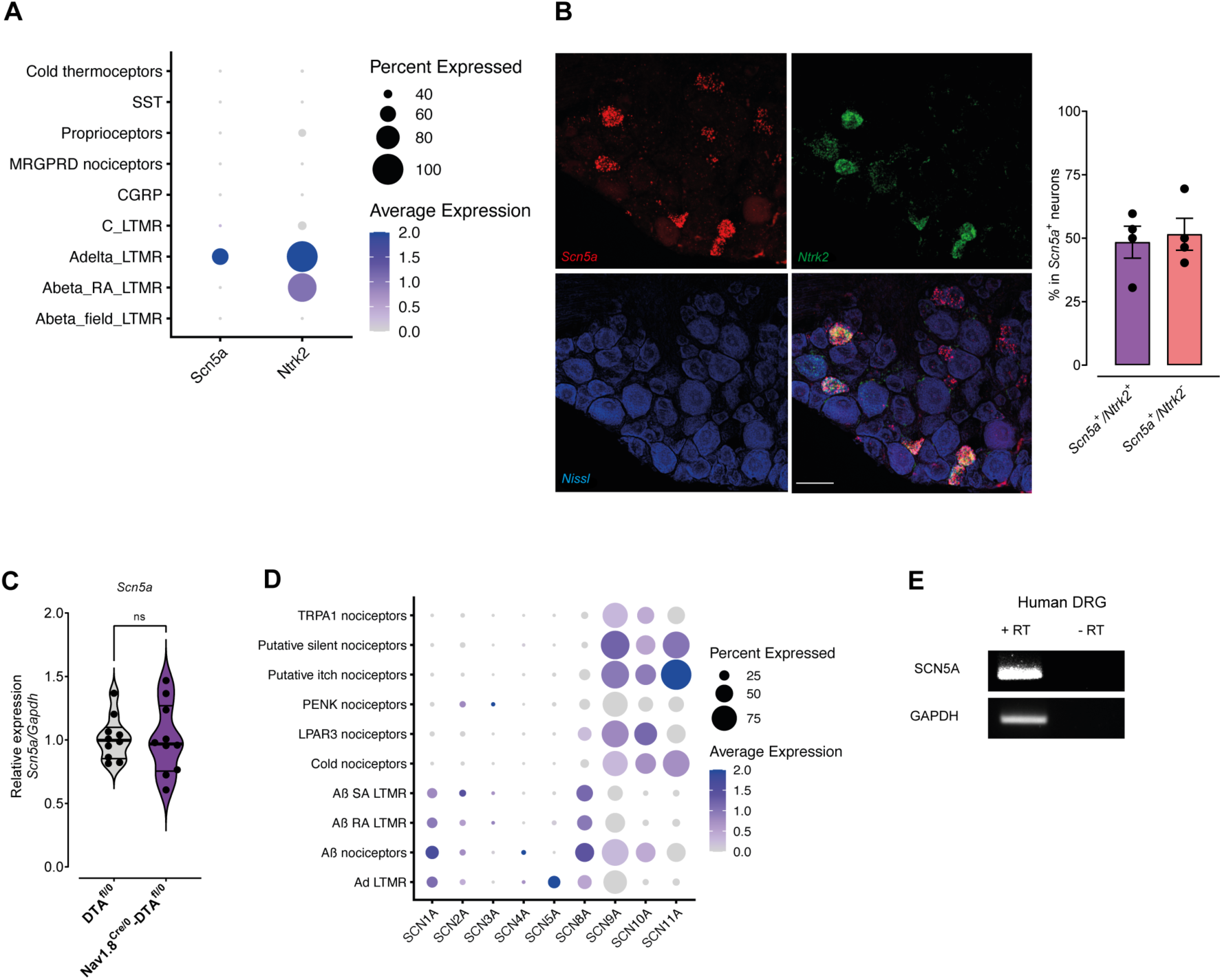
*Scn5a* expression profile in mouse and human sensory ganglia. **(A)** *Scn5a* and *Ntrk2* expression in mouse DRG neurons subpopulations (GSE154659). (**B**) *Scn5a* expression in Ntrk2+ neurons in the mouse DRG by RNA scope (n=4). Scale bar: 50 µm. (**C**) *Scn5a* mRNA expression in the DRGs of mice lacking Nav1.8+ sensory neurons, Student’s test t, *p*>0.05 (n = 10). (**D**) Sodium channels distribution in neuronal subsets of the human DRG (phs001158). (**E**) Validation of *SCN5A* expression in the human DRG by RT-PCR.

We validated these transcriptomic findings anatomically using RNAscope *in situ* hybridization, which confirmed *Scn5a* expression in mouse DRG neurons (14.6 ± 0.8 of total neurons) and demonstrated co-localization with *Ntrk2* (TrkB) (Fig. 3C), confirming its presence in TrkB⁺ neurons sub-populations (Fig. 3C). This expression was corroborated in DRG sections from TrkB-Cre/tdTomato reporter mice (Fig. S2A). In line with the scRNA-seq prediction, *Scn5a*⁺ neurons were predominantly NF200⁺ large-sized neurons (Fig. S2B), and its expression was preserved in DRG samples from Nav1.8-Cre/DTA mice (Fig. 3F), supporting the conclusion that *Scn5a* is enriched in Nav1.8-negative, TrkB⁺ mechanosensory neurons rather than classical Nav1.8⁺ nociceptors.

To assess translational relevance, we examined human DRG spatial single-cell RNAseq datasets (phs001158, Tavares-Ferreira et al., 2022) and found that *SCN5A* expression is restricted to specific neuronal populations, with enrichment in mechanosensory clusters associated with TrkB, mirroring the murine pattern (Fig. 3G). RT-PCR detection of *SCN5A* in human DRG further supports conserved expression across species. Together, these data identify *Scn5a/SCN5A* as a conserved TrkB⁺ mechanosensory neuron-enriched transcript in mouse and human DRG, consistent with recent evidence for Nav1.5 protein in TrkB⁺ sensory neurons (Zhang et al., 2026).

### 4. Paclitaxel-chemotherapy increases Snc5a gene expression in dorsal root ganglia

Nav1.5 shapes action potential initiation and firing in TrkB⁺ mechanosensory neurons, and conditional Nav1.5 deletion in TrkB⁺ sensory neurons reduces mechanical hypersensitivity across pain models (Zhang et al., 2026). Together with our finding that TrkB⁺ fibers drive paclitaxel-evoked mechanical hypersensitivity, this supported testing Nav1.5 in chemotherapy-induced neuropathic pain.

We first assessed *Scn5a* expression in the sciatic nerve, DRG, and spinal cord following paclitaxel treatment (Fig. 4A). Transcriptional analyses revealed significant changes in *Scn5a* expression in tissues from paclitaxel-treated mice compared to naïve animals, as shown by violin plots (Fig. 4B–D). Notably, these changes were restricted to the DRG tissue. Re-analysis of publicly available datasets (Renthal et al., 2022) further indicated that *Scn5a* expression remains highly restricted to NF3/TrkB⁺ DRG neurons (Fig. 4E) and that paclitaxel selectively increases *Scn5a* within this cluster while leaving other neuronal populations, including nociceptor-enriched groups, largely devoid of *Scn5a*. These findings indicate that chemotherapy does not broadly changes *Scn5a* across sensory neurons but instead drives a cell–type–specific upregulation within TrkB⁺ mechanosensory neurons.

**Figure 4.**
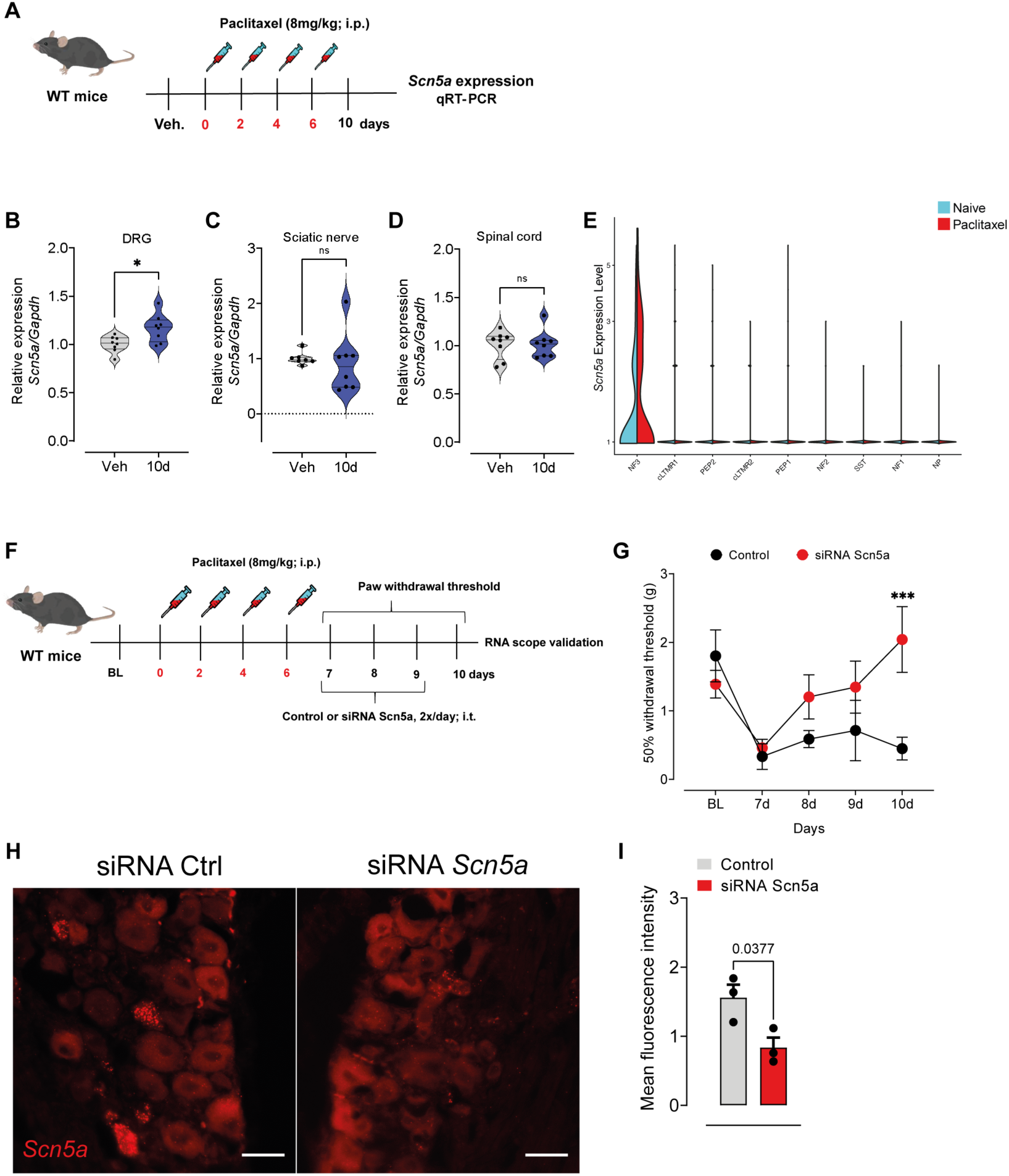
Nav1.5 mediates paclitaxel-induced mechanical pain hypersensitivity. (**A**) Schematic diagram of paclitaxel chemotherapy. (**B-D**) *Scn5a* expression in mice DRG, sciatic nerve and spinal cord after paclitaxel or vehicle treatments, \**p*<0.05 versus vehicle, 1-Way ANOVA, Bonferroni post-test, n= 8-10. (**E**) *Scn5a* expression across neuronal subsets in the mouse DRG after paclitaxel (GSE154659). (**F**) Schematic diagram of the time course of behavioral responses after siRNA against *Scn5a* or control after paclitaxel chemotherapy (8 mg/kg; i.p). (**G**) Mechanical pain hypersensitivity in paclitaxel-treated mice that received siRNA against *Scn5a* or control, \*\*\**p*<0.05 versus control, 2-Way ANOVA, Bonferroni post-test, n =4. (**H**) *In situ* (RNA scope) of *Scn5a* expression in control or *Scn5a* siRNA-treated mice. (**I**) Quantification of mean fluorescence intensity, \**p*<0.05, Student’s t test, n=3.

To determine whether Nav1.5 is functionally required for paclitaxel-induced mechanical hypersensitivity, we performed siRNA-mediated *Scn5a* knockdown via intrathecal delivery (Fig. 4F; Li et al., 2024). At the onset of paclitaxel-induced mechanical pain hypersensitivity, *Scn5a*-targeting siRNA significantly attenuated mechanically evoked pain behaviours compared with control siRNA-treated animals (Fig. 4G). RNA scope analyses of DRG sections confirmed effective *Scn5a* knockdown, as evidenced by reduced transcript signal intensity and decreased mean fluorescence in siRNA-treated mice (Fig. 4H–I). These findings indicate that *Scn5a* is a key contributor to the development of paclitaxel-induced mechanical pain hypersensitivity.

## Conclusion

In summary, our findings identify a previously unrecognized and functionally relevant role for TrkB+ mechanosensory neurons and its molecular expressed entity, Scn5a/Nav1.5, in the development of chemotherapy-induced mechanical pain hypersensitivity. By combining behavioral, transcriptomic, and molecular approaches, we demonstrate that mechanical hypersensitivity induced by paclitaxel is not dependent on classical Nav1.8⁺ nociceptors but instead is critically mediated by TrkB⁺ mechanosensory neurons. Within this population, Scn5a/Nav1.5 emerges as a selectively enriched and dynamically regulated molecular determinant, whose expression is increased following chemotherapeutic exposure and whose silencing is sufficient to attenuate pain-related behaviors.

Importantly, our results are consistent with and extend the recent findings of Zhang et al. (2026), published in *Brain*, which identified Nav1.5 as a key contributor to the electrophysiological properties of TrkB⁺ sensory neurons. While that study provided fundamental insight into the role of Nav1.5 in somatosensory neuron function, our work expands this framework by placing *Scn5a*/Nav1.5 in a clinically relevant context, chemotherapy-induced peripheral neuropathy. From a therapeutic perspective, Nav1.5 is an attractive, mechanistically grounded target because sodium channels are core determinants of afferent excitability and are widely implicated in neuropathic pain pathogenesis. However, Nav1.5 is classically associated with cardiac excitability, raising the possibility that systemic Nav1.5 blockade could carry on-target cardiac liabilities. These liabilities may be mitigated through tissue-targeted strategies, such as the siRNA approach, for which we provide proof of principle and that may have translational potential (Hu et al., 2020).

Together, these findings support a model in which chemotherapy selectively reshapes the molecular landscape of mechanosensory neurons, promoting aberrant excitability through *Scn5a* upregulation and thereby driving mechanical allodynia, a classical feature of neuropathic pain. Taken together, the consistent findings across species, along with the spatial and functional validation, strengthen the evidence for this mechanism and highlight its potential relevance for clinical and therapeutic translation.

## Data availability

The data that support the findings of this study are available on request from the corresponding author.

## Funding

Gomes-Aragão FIF holds a postdoctoral fellowship from the São Paulo Research Foundation (FAPESP) #2024-0499-2, Adjafre BL held a fellowship from the Brazilian National Council for Scientific and Technological Development (CNPq) grant number 201118/2022-0, Lucena-Silva GV holds a FAPESP doctoral fellowship #2021-10494-7, Kanada LG holds a FAPESP fellowship #2025-15219-5, and Cunha TM holds FAPESP grants #2024-10827-4 and #2013-08216-2. Temugin Berta is funded by NIH/NINDS grant NS136108.

## Competing interests

The authors report no competing interests.

## Supplementary material

Supplementary material is available at *Brain* online.

---

**Supplementary Figure 1.**
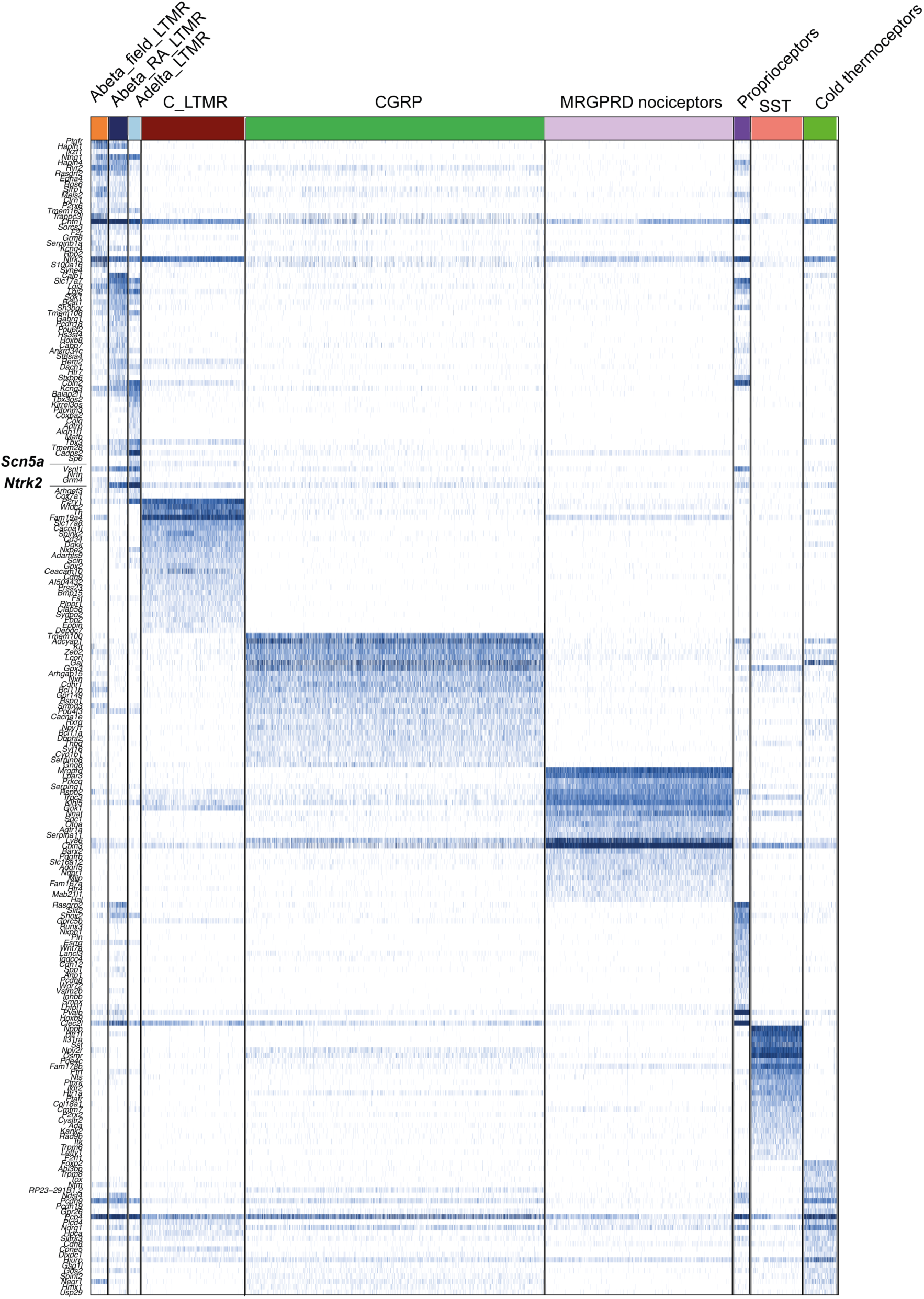
Transcriptomic profile of mouse DRG neurons. Top 25 of differential expressed genes per neuronal subsets (GSE139088).

**Supplementary Figure 2.**
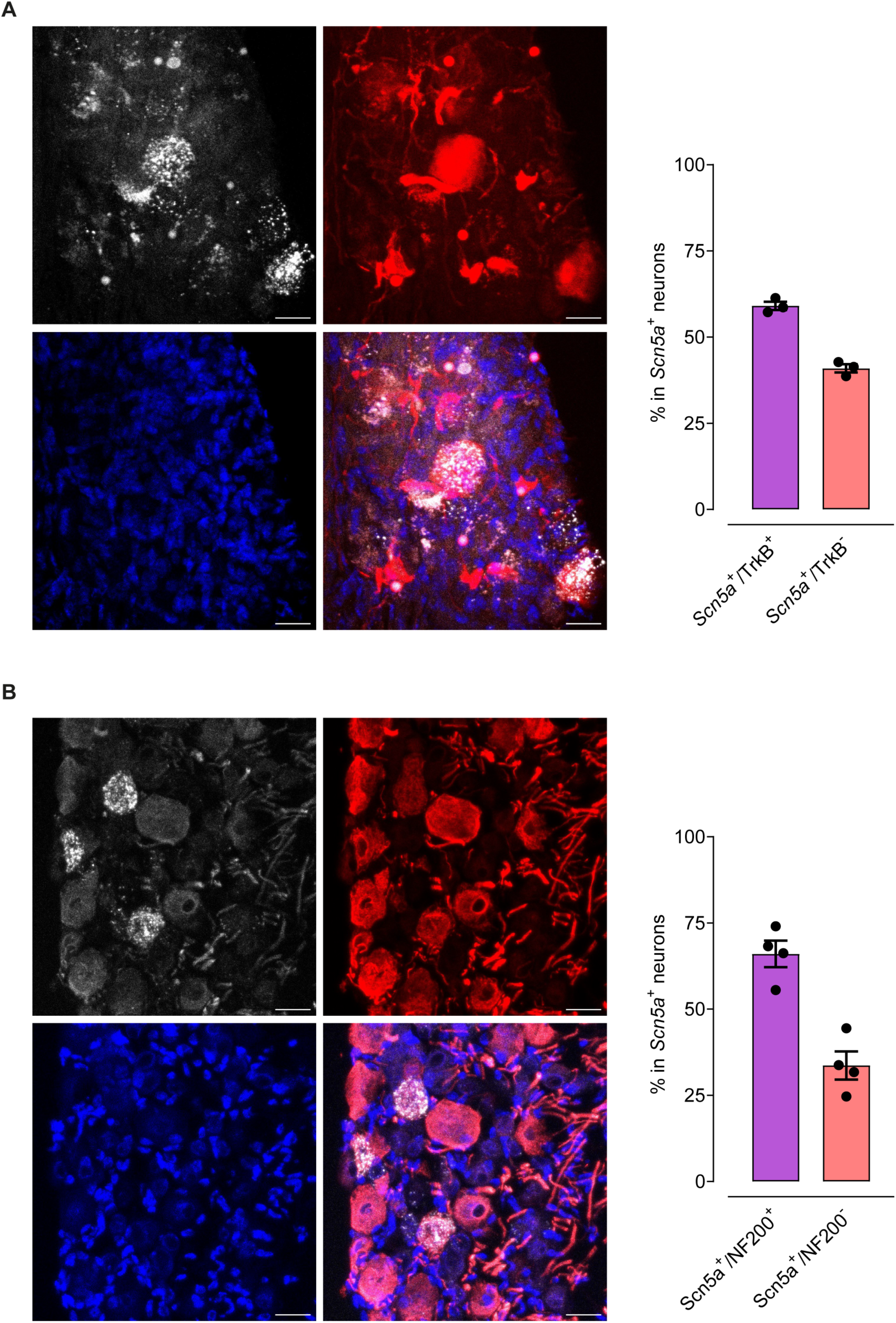
*Scn5a* expression profile in mouse sensory ganglia. (**A**) Presentative images and quantification of *in situ* analyses of *Scn5a* expression in NF200+ (n=4) and (**B**) TrkB-tdTomato+ neurons (n=3) in the mouse DRG. Scale bar: 25µm.

**Table S1.**
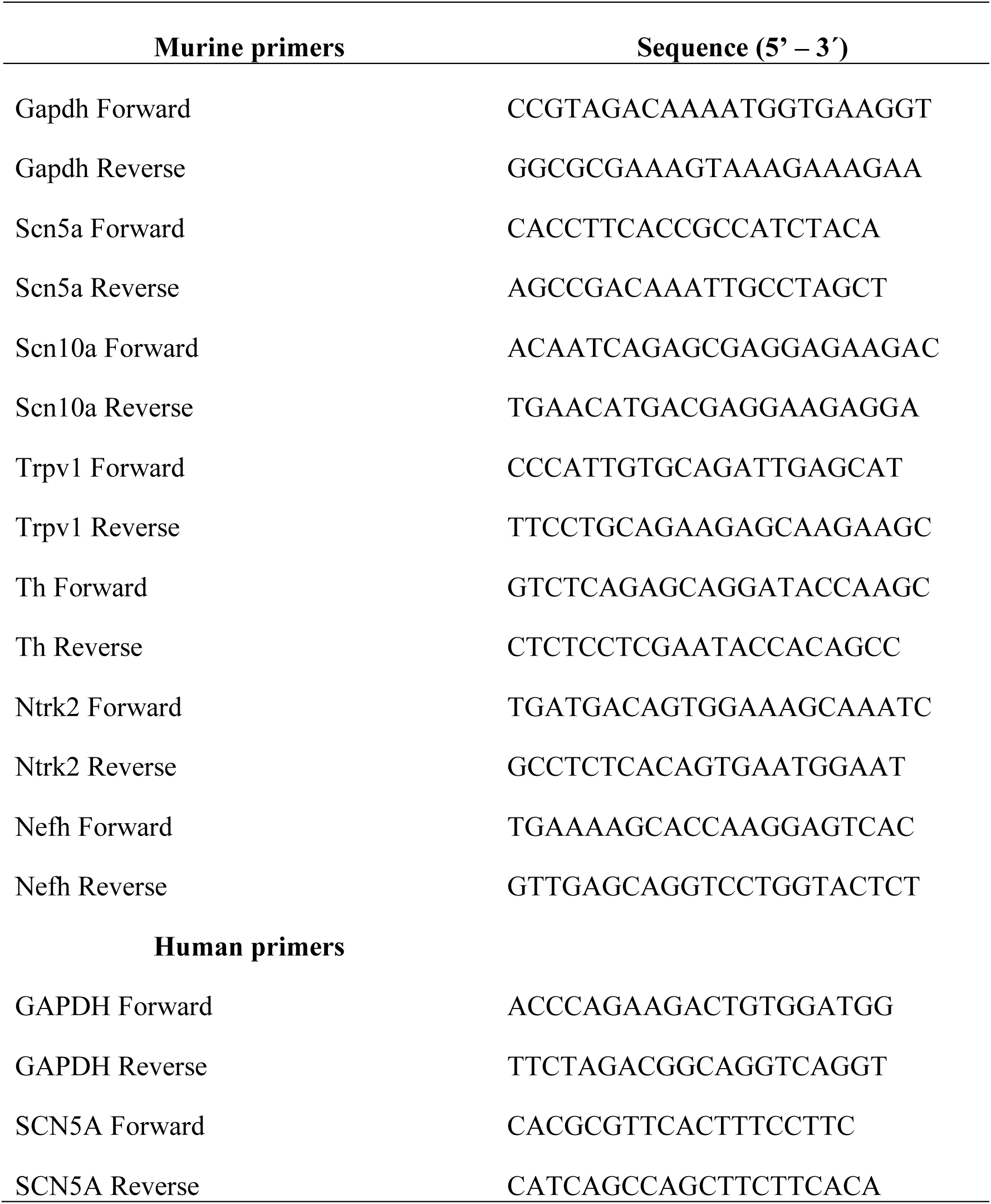
Primers sequence.

## Notes

### Competing Interest Statement

The authors have declared no competing interest.

## References

Abrahamsen B, Zhao J, Asante CO, Cendan CM, Marsh S, Martinez-Barbera JP, Nassar MA, Dickenson AH, Wood JN. The cell and molecular basis of mechanical, cold, and inflammatory pain. Science. 2008 Aug 1;321(5889):702–5. doi: 10.1126/science.1156916. PMID: 18669863.

Bennett DL, et al. The Role of Voltage-Gated Sodium Channels in Pain Signaling. Physiol Rev. 2019 Apr 1;99(2):1079–1151.

Boyette-Davis JA, et al. An updated understanding of the mechanisms involved in chemotherapy-induced neuropathy. Pain Manag. 2018 Sep 1;8(5):363–375.

Chaplan, S.R., Bach, F.W., Pogrel, J.W., Chung, J.M., Yaksh, T.L., 1994. Quantitative assessment of tactile allodynia in the rat paw. J. Neurosci. Methods 53(1): 55–63. doi: 10.1016/0165-0270(94)90144-9.

Cheng L, Duan B, Huang T, Zhang Y, Chen Y, Britz O, Garcia-Campmany L, Ren X, Vong L, Lowell BB, Goulding M, Wang Y, Ma Q. Identification of spinal circuits involved in touch-evoked dynamic mechanical pain. Nat Neurosci. 2017 Jun;20(6):804–814. doi: 10.1038/nn.4549.

Daou I, Beaudry H, Ase AR, Wieskopf JS, Ribeiro-da-Silva A, Mogil JS, Séguéla P. Optogenetic Silencing of Nav1.8-Positive Afferents Alleviates Inflammatory and Neuropathic Pain. eNeuro. 2016 Mar 16;3(1):ENEURO.0140-15.2016. doi: 10.1523/ENEURO.0140-15.2016.

Decosterd I, Woolf CJ. Spared nerve injury: an animal model of persistent peripheral neuropathic pain. Pain. 2000 Aug;87(2):149–158. doi: 10.1016/S0304-3959(00)00276-1.

Dhandapani R, Arokiaraj CM, Taberner FJ, Pacifico P, Raja S, Nocchi L, Portulano C, Franciosa F, Maffei M, Hussain AF, de Castro Reis F, Reymond L, Perlas E, Garcovich S, Barth S, Johnsson K, Lechner SG, Heppenstall PA. Control of mechanical pain hypersensitivity in mice through ligand-targeted photoablation of TrkB-positive sensory neurons. Nat Commun. 2018 Apr 24;9(1):1640. doi: 10.1038/s41467-018-04049-3.

Dixon, W.J., 1980. Efficient analysis of experimental observations. Annu. Rev. Pharmacol. Toxicol. 20: 441–62. doi: 10.1146/annurev.pa.20.040180.002301.

Finnerup NB, et al. Neuropathic Pain: From Mechanisms to Treatment. Physiol Rev. 2021 Jan 1;101(1):259–301.

Fonseca MM, Davoli-Ferreira M, Santa-Cecília F, Guimarães RM, Oliveira FFB, Kusuda R, Ferreira DW, Alves-Filho JC, Cunha FQ, Cunha TM. IL-27 Counteracts Neuropathic Pain Development Through Induction of IL-10. Front Immunol. 2020 Jan 28;10:3059. doi: 10.3389/fimmu.2019.03059.

Fumagalli G, et al. Neuroinflammatory Process Involved in Different Preclinical Models of Chemotherapy-Induced Peripheral Neuropathy. Front Immunol. 2021 Feb 4;11:626687.

Handler A, Ginty DD. The mechanosensory neurons of touch and their mechanisms of activation. Nat Rev Neurosci. 2021 Sep;22(9):521–537. doi: 10.1038/s41583-021-00489-x.

Hao Y, Stuart T, Kowalski MH, Choudhary S, Hoffman P, Hartman A, Srivastava A, Molla G, Madad S, Fernandez-Granda C, Satija R. Dictionary learning for integrative, multimodal and scalable single-cell analysis. Nat Biotechnol. 2024 Feb;42(2):293–304. doi: 10.1038/s41587-023-01767-y.

Hu B, Zhong L, Weng Y, Peng L, Huang Y, Zhao Y, Liang XJ. Therapeutic siRNA: state of the art. Signal Transduct Target Ther. 2020 Jun 19;5(1):101. doi: 10.1038/s41392-020-0207-x.

Hylden JL, Wilcox GL. Intrathecal morphine in mice: a new technique. Eur J Pharmacol. 1980;67(2-3):313–6.

Li X, Prudente AS, Prato V, Guo X, Hao H, Jones F, Figoli S, Mullen P, Wang Y, Tonello R, Lee SH, Shah S, Maffei B, Berta T, Du X, Gamper N. Peripheral gating of mechanosensation by glial diazepam binding inhibitor. J Clin Invest. 2024 Jun 18;134(16):e176227. doi: 10.1172/JCI176227.

Luiz AP, MacDonald DI, Santana-Varela S, Millet Q, Sikandar S, Wood JN, Emery EC. Cold sensing by NaV1.8-positive and NaV1.8-negative sensory neurons. Proc Natl Acad Sci U S A. 2019 Feb 26;116(9):3811–3816. doi: 10.1073/pnas.1814545116.

McDougall JJ, O’Brien MS. Analgesic potential of voltage gated sodium channel modulators for the management of pain. Curr Opin Pharmacol. 2024 Jan 25;75:102433.

Percie du Sert N, Hurst V, Ahluwalia A, Alam S, Avey MT, Baker M, Browne WJ, Clark A, Cuthill IC, Dirnagl U, Emerson M, Garner P, Holgate ST, Howells DW, Karp NA, Lazic SE, Lidster K, MacCallum CJ, Macleod M, Pearl EJ, Petersen OH, Rawle F, Reynolds P, Rooney K, Sena ES, Silberberg SD, Steckler T, Würbel H. The ARRIVE guidelines 2.0: Updated guidelines for reporting animal research. Br J Pharmacol. 2020 Aug;177(16):3617–3624. doi: 10.1111/bph.15193.

Renganathan M, Dib-Hajj S, Waxman SG. Na(v)1.5 underlies the ‘third TTX-R sodium current’ in rat small DRG neurons. Brain Res Mol Brain Res. 2002 Oct 15;106(1-2):70–82. doi: 10.1016/s0169-328x(02)00411-4.

Renthal W, Tochitsky I, Yang L, Cheng YC, Li E, Kawaguchi R, Geschwind DH, Woolf CJ. Transcriptional Reprogramming of Distinct Peripheral Sensory Neuron Subtypes after Axonal Injury. Neuron. 2020 Oct 14;108(1):128–144.e9. doi: 10.1016/j.neuron.2020.07.026.

Saika F, Matsuzaki S, Kobayashi D, Ideguchi Y, Nakamura TY, Kishioka S, Kiguchi N. Chemogenetic Regulation of CX3CR1-Expressing Microglia Using Gi-DREADD Exerts Sex-Dependent Anti-Allodynic Effects in Mouse Models of Neuropathic Pain. Front Pharmacol. 2020 Jun 19;11:925. doi: 10.3389/fphar.2020.00925.

Santana-Varela S, Bogdanov YD, Gossage SJ, Okorokov AL, Li S, de Clauser L, Alves-Simoes M, Sexton JE, Iseppon F, Luiz AP, Zhao J, Wood JN, Cox JJ. Tools for analysis and conditional deletion of subsets of sensory neurons. Wellcome Open Res. 2021 Sep 30;6:250. doi: 10.12688/wellcomeopenres.17090.1.

Schindelin, J., Arganda-Carreras, I., Frise, E., Kaynig, V., Longair, M., Pietzsch, T., Preibisch, S., Rueden, C., Saalfeld, S., Schmid, B., Tinevez, J.Y., White, D.J., Hartenstein, V., Eliceiri, K., Tomancak, P., Cardona, A., 2012. Fiji: an open-source platform for biological-image analysis. Nat. Methods 9(7): 676–82. doi: 10.1038/nmeth.2019.

Schmittgen TD, Livak KJ. Analyzing real-time PCR data by the comparative C(T) method. Nat Protoc. 2008;3(6):1101–8. doi: 10.1038/nprot.2008.73. PMID: 18546601.

Seretny M, et al. Incidence, prevalence, and predictors of chemotherapy-induced peripheral neuropathy: A systematic review and meta-analysis. Pain. 2014 Dec;155(12):2461–2470.

Sharma N, Flaherty K, Lezgiyeva K, Wagner DE, Klein AM, Ginty DD. The emergence of transcriptional identity in somatosensory neurons. Nature. 2020 Jan;577(7790):392–398. doi: 10.1038/s41586-019-1900-1. Epub 2020 Jan 8. Erratum in: Nature. 2025 Nov;647(8091):E6. doi: 10.1038/s41586-025-09817-y.

Silva NR, Gomes FIF, Lopes AHP, Cortez IL, Dos Santos JC, Silva CEA, Mechoulam R, Gomes FV, Cunha TM, Guimarães FS. The Cannabidiol Analog PECS-101 Prevents Chemotherapy-Induced Neuropathic Pain via PPARγ Receptors. Neurotherapeutics. 2022 Jan;19(1):434–449. doi: 10.1007/s13311-021-01164-w.

Tavares-Ferreira D, Shiers S, Ray PR, Wangzhou A, Jeevakumar V, Sankaranarayanan I, Cervantes AM, Reese JC, Chamessian A, Copits BA, Dougherty PM, Gereau RW 4th, Burton MD, Dussor G, Price TJ. Spatial transcriptomics of dorsal root ganglia identifies molecular signatures of human nociceptors. Sci Transl Med. 2022 Feb 16;14(632):eabj8186. doi: 10.1126/scitranslmed.abj8186.

Yang D, Jacobson A, Meerschaert KA, Sifakis JJ, Wu M, Chen X, Yang T, Zhou Y, Anekal PV, Rucker RA, Sharma D, Sontheimer-Phelps A, Wu GS, Deng L, Anderson MD, Choi S, Neel D, Lee N, Kasper DL, Jabri B, Huh JR, Johansson M, Thiagarajah JR, Riesenfeld SJ, Chiu IM. Nociceptor neurons direct goblet cells via a CGRP-RAMP1 axis to drive mucus production and gut barrier protection. Cell. 2022 Oct 27;185(22):4190–4205.e25. doi: 10.1016/j.cell.2022.09.024. Epub 2022 Oct 14.

Zhang CJ, Ji MJ, Zhou XL, Li S, Xu PF, Wu H, Chen Q, Zhao J, Chen XZ, Cox JJ, Zhou XL. Nav1.5 in the dorsal root ganglion plays a crucial role in mechanical hypersensitivity. Brain. 2026 Mar 11:awaf438.

Zhu H, Aryal DK, Olsen RH, Urban DJ, Swearingen A, Forbes S, Roth BL, Hochgeschwender U. Cre-dependent DREADD (Designer Receptors Exclusively Activated by Designer Drugs) mice. Genesis. 2016 Aug;54(8):439–46. doi: 10.1002/dvg.22949. Epub 2016 Jun 3. PMID: 27194399; PMCID: PMC4990490.

